# Frequent coexistence of closely related tree species on a small scale in temperate evergreen forests in Japan

**DOI:** 10.1101/2021.02.03.429543

**Authors:** Shuntaro Watanabe, Yuri Maesako

## Abstract

Understanding how biotic interaction affects species composition and distribution is a major challenge in community ecology. In plants, negative reproductive interaction among closely related species (i.e., reproductive interference) is known to hamper the coexistence of congenic species. Since the magnitude of reproductive interference in plants depends on pollen flow distance, we hypothesized that the coexistence of congeners on a small spatial scale would be less likely to occur by chance but that such coexistence would be likely to occur on a scale larger than pollen flow distance. In the present study, we tested this hypothesis using spatially explicit woody plant survey data. Contrary to our prediction, congenic tree species often coexisted at the finest spatial scale and significant exclusive distribution was not detected. Our results suggest that cooccurrence of congenic tree species is not structured by reproductive interference, and they indicate the need for further research to explore the factors that mitigate the effects of reproductive interference.

## Introduction

Understanding how biotic interaction affects species composition and distribution is a major ongoing challenge in community ecology. Among biotic interactions, competition is the most important and well-studied interaction (Goldberg and Barton 1992). A common hypothesis related to the role of competition in community assembly, termed the competition-relatedness hypothesis (CRH; Cahill et al. 2008), states that closely related species compete more intensely than distantly related species, which hypothetically limits the ability of closely related species to coexist (Webb et al. 2002; Slingsby and Verboom 2006; Prinzing et al. 2008; reviewed by Mayfield and Levine 2010; HilleRisLambers et al. 2012).

The findings of Elton (1946), i.e., that a lower number of species per genus are observed in local areas than in the entire United Kingdom, appear to provide evidence for the competitive exclusion of ecologically similar congeners in local habitats. Although the CRH has been widely discussed (Dayan and Simberloff 2005), empirical support for the hypothesis remains inconclusive (Webb et al. 2002; Cavender-Valles et al. 2004; reviewed by Mayfield and Levine 2010). In one study, Cahill et al. (2008) tested the CRH in plant species using experimental data; they revealed that the relationships between phylogenetic relatedness and competitive ability differed between monocots and eudicots.

Although the community phylogenetic evidence for the CRH in plants is mixed, recent research has shown that negative reproductive interaction among closely related species (i.e., reproductive interference) limits the coexistence of closely related species. Reproductive interference is defined in the present study as the negative fitness effects of pollen flow among species. Such effects may be pre- or post-zygotic, and they directly affect reproductive success through responses such as reduced fertilization due to pollen tube competition or the production of unviable hybrids (e.g., Brown and Mitchell 2001; Brown et al. 2002; Takakura et al. 2009; 2011; Takakura and Fujii 2010; Nishida et al. 2014). Given that reproductive interference involves positive frequency-dependent selection (Kuno 1992), it can rapidly lead to extinction of the affected species with lower population density (Kishi and Nakazawa 2012).

Previous studies of reproductive interference have provided some insight into the pattern and spatial scale of closely related species’ coexistence. Because shared recent ancestry can yield shared reproductive traits (including similarities in the timing of reproduction, mate recognition, pollination system, and gamete recognition) close relatives (e.g., congeners and sister taxa) are less likely to coexist by chance on a local scale (Whitton et al. 2018). Additionally, the extent of reproductive interference in plants depends on pollen flow distance (Takakura et al. 2009). Therefore, it can be hypothesized that the coexistence of congeners on a small spatial scale is less likely to occur by chance, whereas coexistence is more likely to occur on scale larger than pollen flow distance.

In the present study, we aimed to quantitatively assess the distribution patterns of closely related woody plant species in the native forests of Japan to determine the effects of species interactions, especially reproductive interference, on forest community assembly. Using spatially explicit woody plant survey data, we tested the following predictions: (1) congenic species do not coexist on a fine spatial scale but coexist on large spatial scale; and (2) on a large spatial scale, congenic species show an exclusive distribution.

## Materials and method

### Study site

The study area (~1 km^2^) was located in the Kasugayama primary forest, Nara prefecture, western Japan (34’41’N 135’51’E) (Fig. 1). Because the forest has been preserved as a holy site of the Kasuga Taisha shrine, hunting and logging have been prohibited there since 841 AD (Maesako et al. 2007). In the area, the mean annual temperature in 2019 is 16.3°C and the average annual precipitation in 2019 is 1482.5 mm. The highest point of the forest is 498 m. The natural vegetation in the area is evergreen broadleaved forest (Naka 1982); however, the deer population has recently increased in the forest, causing the spread of alien species such as *Sapium sebiferum* and *Nagia nagi* (Maesako et al. 2007).

**Fig. 1.**
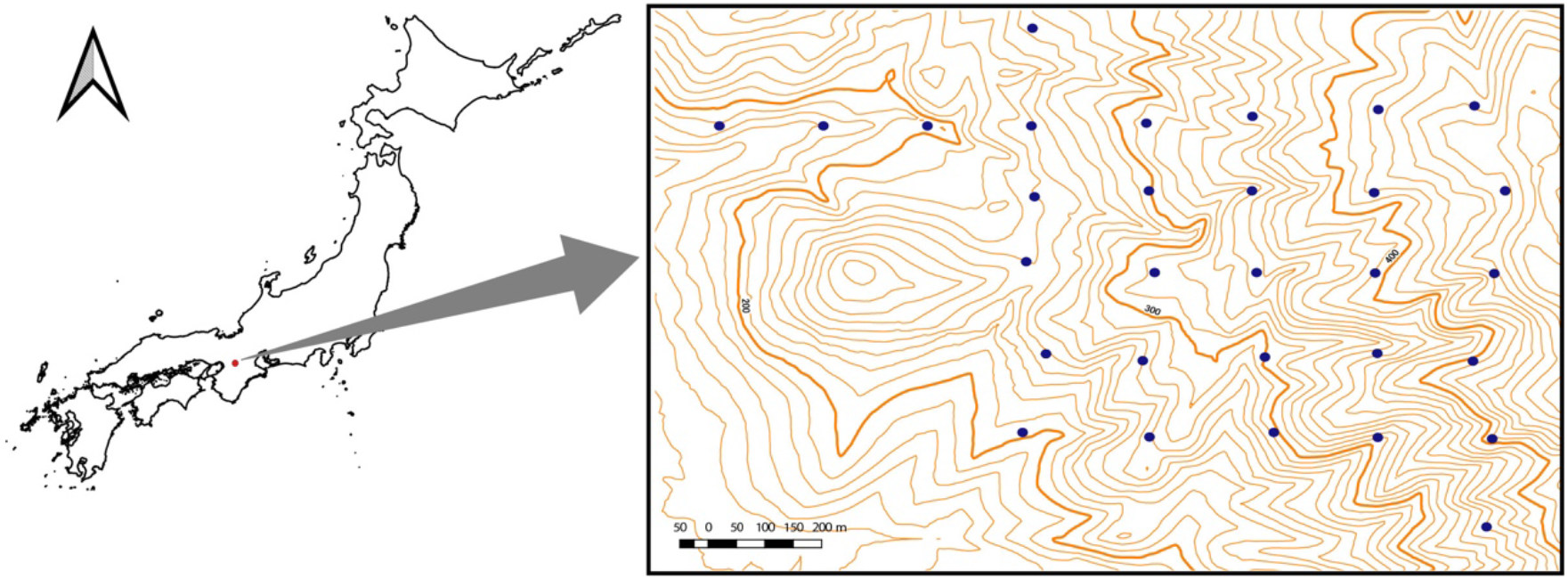
Location of the study site (~1 km^2^) at Kasugayama primary forest, Nara prefecture, western Japan. The specific locations of the study plots are denoted by dots. Scale bar: 200 m.

### Field survey

Field studies were conducted from June to September 2015. In the study area, 30 transect plots (~0.1 ha in size) were established (Fig. 1). Tree species richness was surveyed in each plot; all tree species with heights >130 cm were recorded. Species were grouped into three life-form categories, namely trees, shrubs, or lianas, following Satake (1981–1982). For categorization of small plants, species with a diameter at breast height (DBH) >10 cm on average were classified as trees while species with a DBH <10 cm were classified as shrubs.

### Interspecific competition and null model analysis

We employed the species-to-genus ratio (S/G ratio) as an indicator of intragenic interactions for the categorized tree, shrub, and liana species. The S/G ratio has long been used to describe community patterns and to infer levels of competitive interactions among species within genera (reviewed by Simberloff 1970). A low S/G ratio can be interpreted as a product of strong intrageneric competition (Elton 1946), which could limit congeneric coexistence (Darwin 1859). First, we calculated an S/G ratio for the whole area and then tested the deviation of this S/G ratio from a 1:1 ratio using a z-test. Second, to test spatial scale dependencies, we calculated S/G ratios at five *a priori*-defined spatial scales (0.1, 4, 16, 36, and 64 ha).

### Distribution exclusiveness among congener species

To evaluate the exclusivity of congeners, we calculated the checkerboard scores (C-scores; Stone and Roberts 1990) for the genera with multiple species of the same genus distributed within the study area and that occurred in more than three plots (*Quercus*, *Carpinus*, and *Prunus*). Note that there are divisions of opinion among the researchers on the question that should genus *Prunus* treat as a single genus or not (Ohba 1992; Mabberley 2008). Therefore, the genus *Prunus* in this study includes the subgenera *Cersus* and *Laurocerasus*.

We set *r*_*i*_ and *r*_*j*_ as the number of plots in which species *i* and *j*, respectively, were present; the checker unit *C_ij_* associated with the two species was defined as follows:

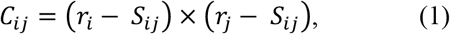

where *S*_*ij*_ indicates the extent of copresence (i.e., the number of plots shared by the two species).

For *N* species, there are *P* = *N*(*N* − 1)/2 species pairs; thus, the C-score is calculated as follows:

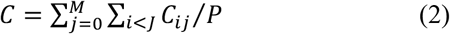

The C-score becomes larger as the two species occur more commonly in different plots. We simulated null models to compare the observed C-score with stochastic distributions. The null models, which were run 999 times for each species pair, randomly shuffle the number of species (α-diversity) among sampling locations while preserving the number of species in the species pool (γ-diversity). All statistical analyses were performed using R software version 3.6.1 (R Core Team 2019). We used the package EcoSimR (Gotelli et al. 2015) to compute C-score.

## Results

### S/G ratio

At the study site, we recorded 42 tree species from 31 genera, 20 shrub species from 19 genera, and seven liana species from six genera. The resultant S/G ratios for trees, shrubs, and lianas were 1.350, 1.021, and 1.200, respectively. Only the S/G ratio for tree species significantly deviated from the 1:1 ratio, whereas those for shrub and liana species did not (Fig. 2).

**Fig. 2.**
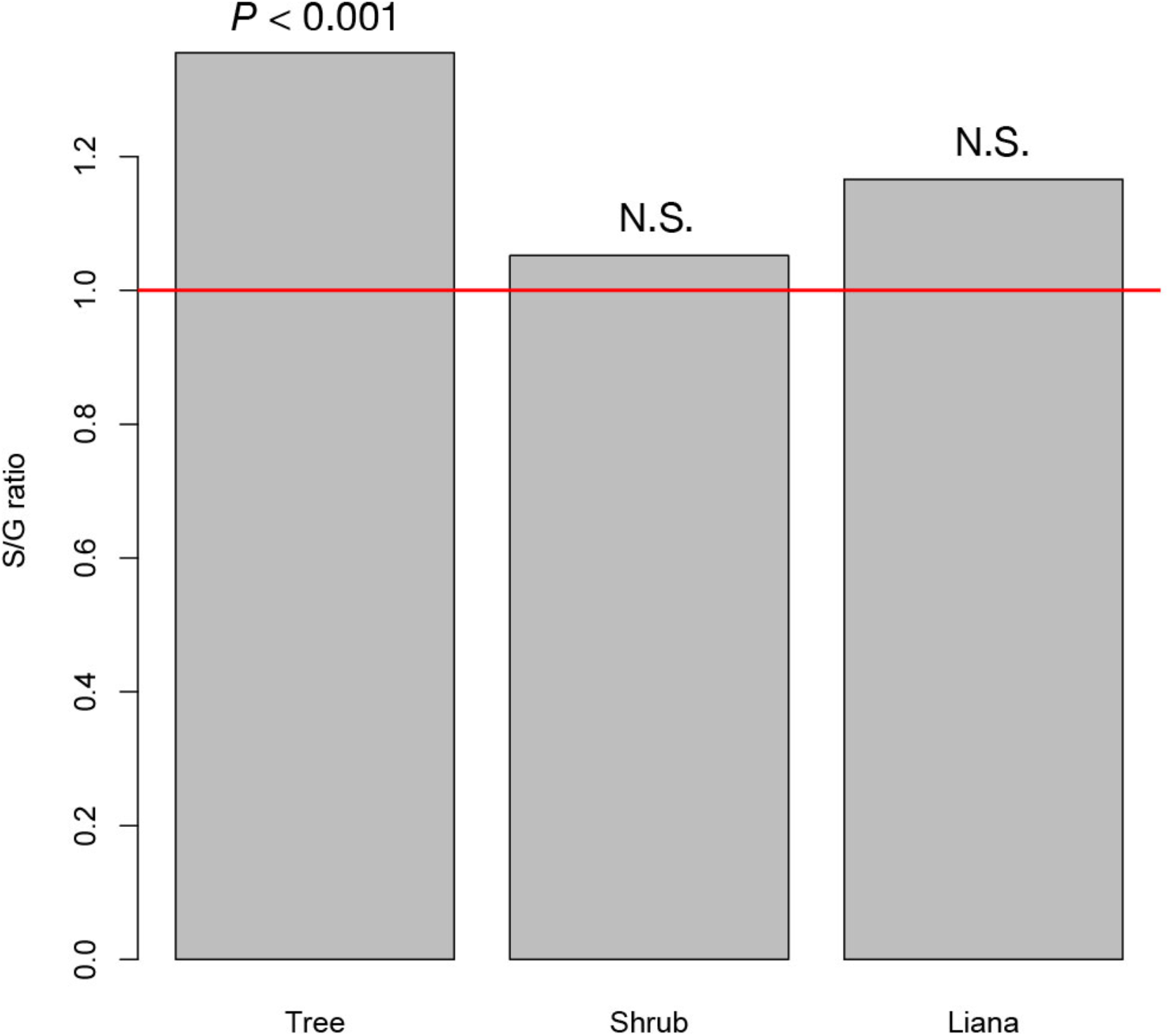
The species–to-genus ratios for the tree, shrub, and liana species categorized in the study site. This ratio significantly deviated from the 1:1 ratio (red horizontal line) for tree species but not shrub or liana species.

The average S/G for trees increased as spatial scale increased. Even at the smallest spatial scale, however, the average S/G ratio for trees exceeded the 1:1 ratio (Fig. 3), indicating that the coexistence of congeners frequently occurs at the smallest spatial scale for tree species.

**Fig. 3.**
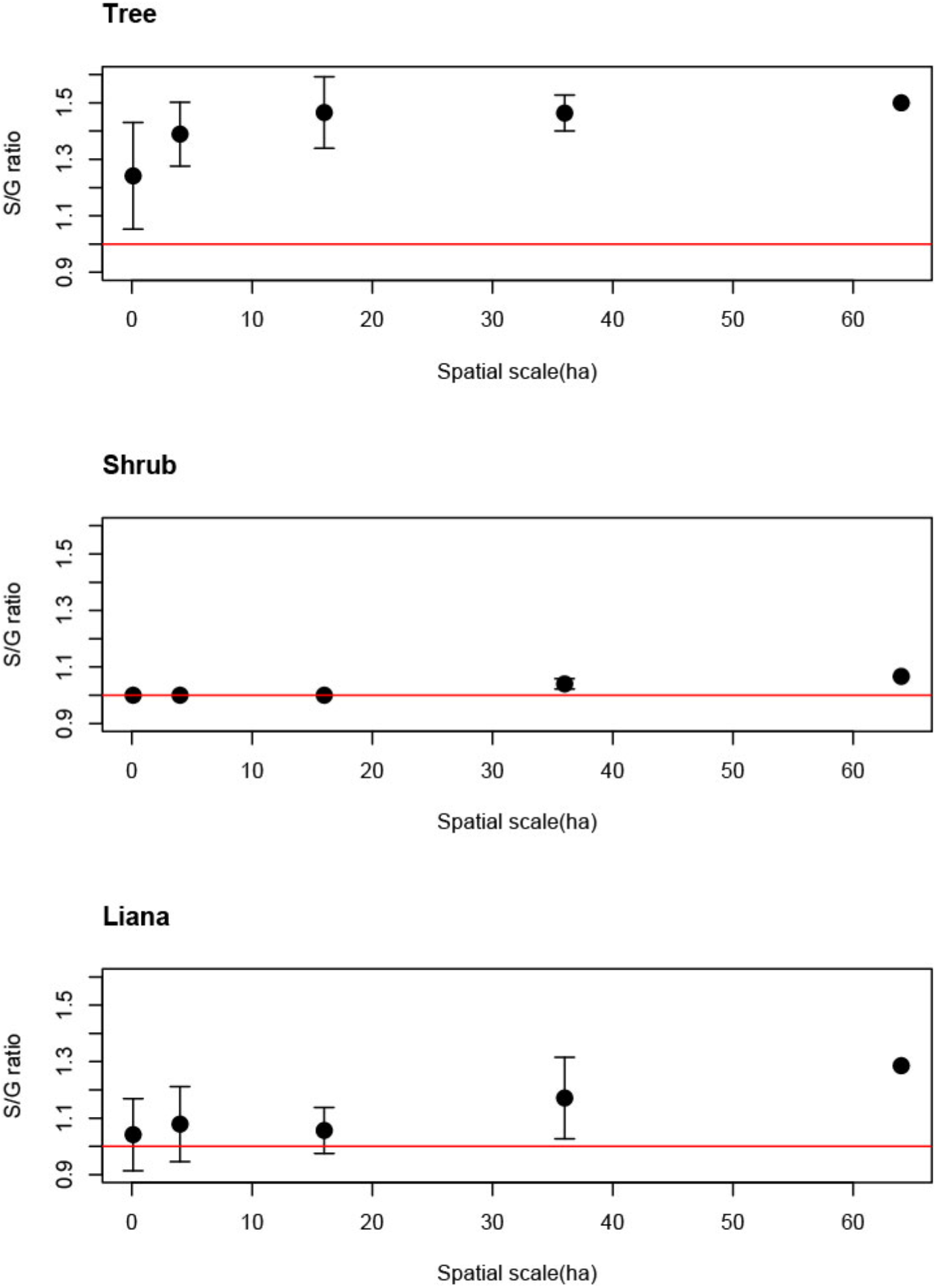
Spatial scale dependencies of species-to-genus ratios for the tree, shrub, and liana species categorized in the study site. Error bars represent standard deviations. Red lines indicate the 1:1 ratio.

### C- score

The C-score did not fall outside the 95% confidence intervals of the null model distribution for all genera, indicating that statistically significant exclusive distribution of species from the same genus did not occur (Fig. 4).

**Fig. 4.**
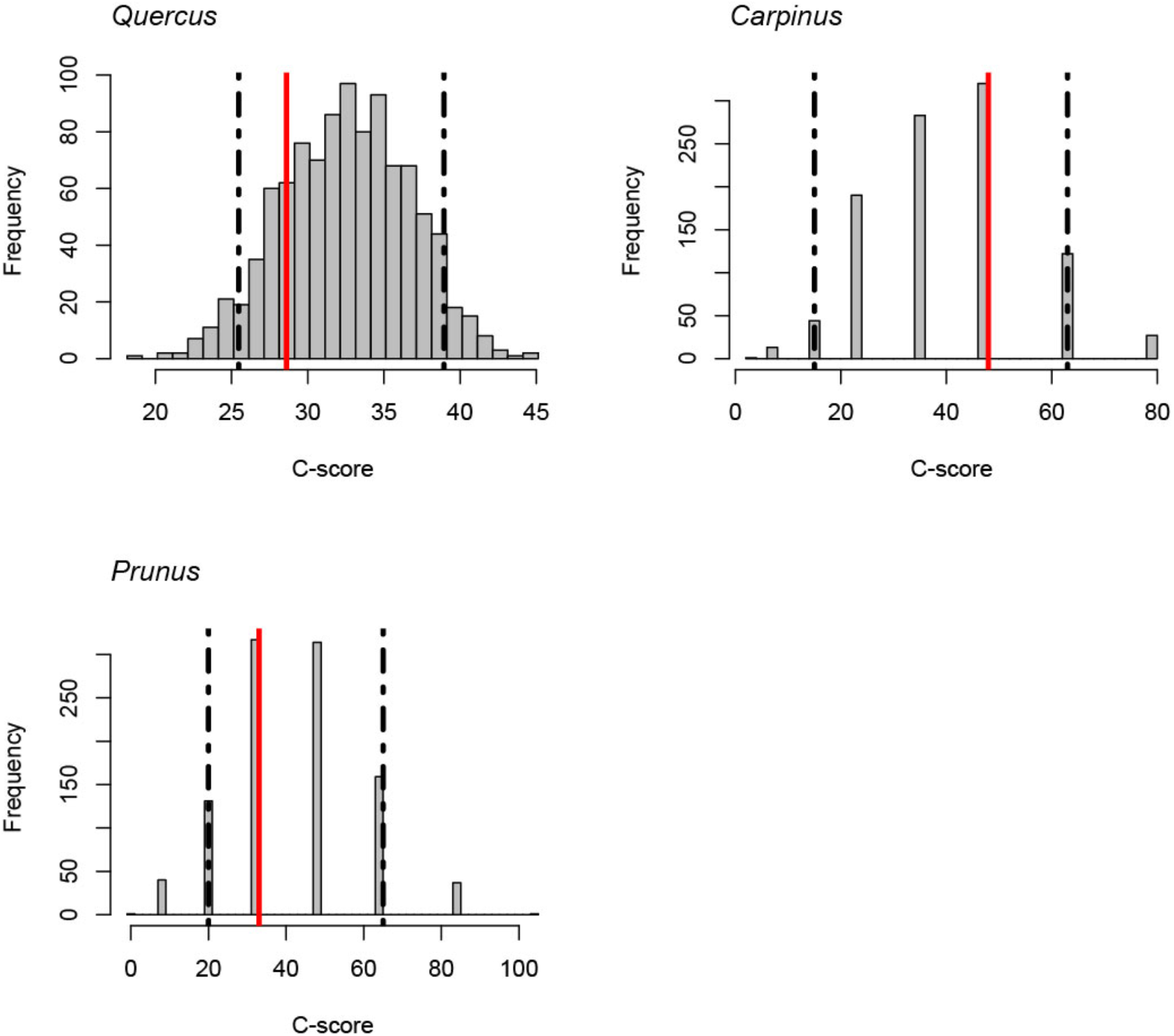
C-scores (checkerboard score; Stone and Roberts 1990) for the genera *Quercus*, *Carpinus* and *Prunus*. Red lines indicate the observed C-score. Histograms indicate the null model distribution. Broken lines indicate the 95% (upper and lower) confidence intervals of the null model distribution.

## Discussion

Our results show that, at least in our study area, closely related tree species often coexist even at the finest spatial scale and that statistically significant exclusive distribution of species from the same genus does not occur, suggesting that cooccurrence of congenic tree species is not structured by reproductive interference. Previous theoretical studies predicted that reproductive interference could more readily prevent coexistence when compared to resource competition (Kuno 1992; Kishi and Nakazawa 2013). Additionally, empirical studies have demonstrated that reproductive interference can cause rapid congenic species exclusion (Takakura et al. 2009; Takakura and Fujii 2010; Runquist and Stanton 2012). Thus, mitigation of reproductive interference is likely required to maintain coexistence of closely related species.

In plants, the spatial extent of reproductive interference corresponds to pollen flow distance (Takakura et al. 2011); consequently, coexistence of closely related plant species is expected at the spatial extent to which pollen flow does not occur. However, the frequent pollen dispersal range of the dominant tree genera in our study site (i.e., *Quecus*, *Acer*, and *Machilus*) is within a few tens of meters (Nakanishi et al. 2004; Kikuchi et al. 2009; Watanabe et al. 2018). This suggests that the congenic tree species in our study site coexist on spatial scales at which reproductive interference can occur frequently and that, given the spatial factor, the effects of reproductive interference could not be avoided in this study. A previous theoretical study suggested that recruitment fluctuation could enable coexistence of closely related tree species on a local scale by producing temporal resource partitioning (a mechanism known as the storage effect) (Usinowicz et al. 2012). One of the dominant genera in our study area, *Quecus*, shows considerable variation in annual seed production (Hirayama et al. 2012), which might contribute to maintaining the coexistence of congener species. Another mechanism that could potentially weaken reproductive interference is reproductive character displacement (Pfennig and Pfennig 2009). Eaton et al. (2012) showed that disparity in the floral traits of plants could reduce negative reproductive interactions among closely related species. However, there remains a lack of direct evidence to show that reproductive character displacement reduces the effect of reproductive interference. Moreover, experimental evidence of reproductive interference is limited to herbaceous plants. Therefore, the mechanism underlying mitigation of reproductive interference during tree community assembly requires further investigation.

In the present study, the S/G ratio for shrubs and lianas did not deviate significantly from the 1:1 ratio. This indicates that few congenic shrub or liana species are distributed in the study area (~1 km^2^). Since a low S/G ratio is generally a product of strong intrageneric competition (Elton 1946), this result also suggests the existence of competitive exclusion in shrub and liana species. However, the S/G ratio depends on the number of species present, and it would be expected to decrease in small communities regardless of competition levels (Gotelli and Colwell 2001). In the study area, the deer population has recently increased, and this has affected the regeneration process and species richness of plant species (Shimoda et al. 1994; Maesako 2015). As shrubs are likely to be more susceptible to deer grazing, a reduction in the number of shrub species due to deer grazing may have reduced the shrub S/G ratio. Nevertheless, it should be noted that closely related species of *Neolitsea aciculate*, a dominant shrub in our study area, are distributed about 15 km from the study site (Murata 1977). This suggests that the low S/G ratio for shrubs and lianas would likely increase if the study area were to be extended by tens of kilometers.

In future research, investigation on a larger spatial scale will be required to determine the relationship between plant life history and the spatial scale of exclusive distribution. Previous studies on herbaceous plants suggest that reproductive interference plays an important role in community assembly (Eaton et al 2012; Whitton et al 2018); however, our results, in concert with prior studies, indicate that reproductive interference is somewhat less effective than expected, especially in long-lived plant species. Further study is therefore necessary to identify the key life history traits that mitigate the effects of reproductive interference.

## Acknowledgments

We thank Kohmei Kadowaki for helpful discussion. We also thank Tomoya Inada, Minori Hikichi, and Kayo Takasu for fieldwork assistance. The investigation in this study was permitted by Nara Park Management Office.

